# Intraflagellar transport proteins undergo nonaxonemal staged hindrance between the recruiting distal appendages and the cilium

**DOI:** 10.1101/227587

**Authors:** T. Tony Yang, Minh Nguyet Thi Tran, Weng Man Chong, Chia-En Huang, Jung-Chi Liao

## Abstract

Primary cilia play a vital role in cellular sensing and signaling [1]. An essential component of ciliogenesis is intraflagellar transport (IFT), which first requires IFT-protein recruitment, IFT-protein–motor-protein assembly, axonemal engagement of IFT-protein complexes, and transition zone (TZ) gating [2–9]. The mechanistic understanding of these processes at the ciliary base was largely missing, because it is exceedingly challenging to observe the motion of IFT proteins in this crowded region using conventional microscopy. Here, we report short trajectory tracking of IFT proteins at the base of mammalian primary cilia by optimizing single-particle tracking photoactivated localization microscopy (sptPALM) [10, 11], balancing the imaging requirements of tracking speed, tracking duration, and localization precision for IFT88-mEOS4b in live human retinal pigment epithelial (hTERT-RPE-1) cells. Intriguingly, we found that mobile IFT proteins “switched gears” multiple times from the distal appendages (DAPs) to the ciliary compartment (CC), moving slowly in the DAPs, relatively fast in the proximal TZ, slowly again in the distal TZ, and then much faster in the CC. They could travel through the space between the DAPs and the axoneme without following DAP structures, and reached the space enveloped by the ciliary pocket in the proximal TZ. Together, our live-cell superresolution imaging revealed region-dependent slowdown of IFT proteins at the ciliary base, shedding light on staged control of ciliogenesis homeostasis.

## RESULTS AND DISCUSSION

### Single particle tracking of superresolution localization microscopy enabled dynamic studies of IFT proteins

We developed a single-particle tracking photoactivated localization microscopy (sptPALM)-based imaging protocol using photoconvertible fluorescent proteins to explore the dynamics of intraflagellar transport (IFT) proteins at the base of primary cilia. The excessive crowding of IFT proteins in this region and their high velocities necessitated multi-parameter optimization of sptPALM to balance signal-to-noise ratio (SNR), frame rate, trajectory duration, phototoxicity, and emission isolation. A human retinal pigment epithelial (RPE-1) cell line with stably expressed IFT88 fused to mEos4b, a monomeric photoconvertible protein [12], was created. The cell morphology, average ciliary length, and ciliation frequency of this cell line were similar to those of wild-type RPE-1 cells. Single-molecule detection was achieved by implementing 405-nm photoconversion with its power being gradually and carefully incremented to photoconvert a finite population of IFT88-mEos4b fluorophores that can be optically isolated in this highly crowded region. These photoconverted fluorophores were then excited by a 561-nm laser (Figure 1A, Movie S1). The laser power was optimized such that it had to be strong enough to reach an acceptable SNR while weak enough to extend the imaging duration to be as long as possible and weak enough to minimize phototoxicity. Similar to IFT88 in wild-type RPE-1 cells, IFT88-mEos4b also formed a high-intensity punctum at the ciliary base (Figure 1B). In contrast to most other sptPALM-based studies, which collect data from a cellular or subcellular volume of about 1000 μm^3^, our imaging volume was usually confined to less than 1 μm^3^, and thus the number of emitting events in each experimental run was very limited. Each of the photoconverted mEos4b emitting events was recorded as a short trajectory of IFT88 movement lasting several to tens of time points (Figure 1A). Imaging at 30 ms per frame was adopted to collect enough photons for single molecule fitting while being fast enough to trace IFT88 fast diffusing movement. Each experimental run lasted about 10,000 frames before prevailing photobleaching. Cilia with obvious twitching or shifting motion were excluded from the analysis to avoid its effect on IFT88 dynamics (Figure S1). Simulated experimental runs were carried out to assure our displacement and velocity analyses were valid under the conditions of the sampling rate and the resolution of our experiments (Figure S2). We have examined the effect of pH and found that under the laser power we used (see *Methods)*, our standard cell-culture medium, whose pH is 7.3 to 7.6, is close to optimal for the density and duration of the fluorophores at the base of primary cilia (Figures S3 and S4).

**Figure 1.**
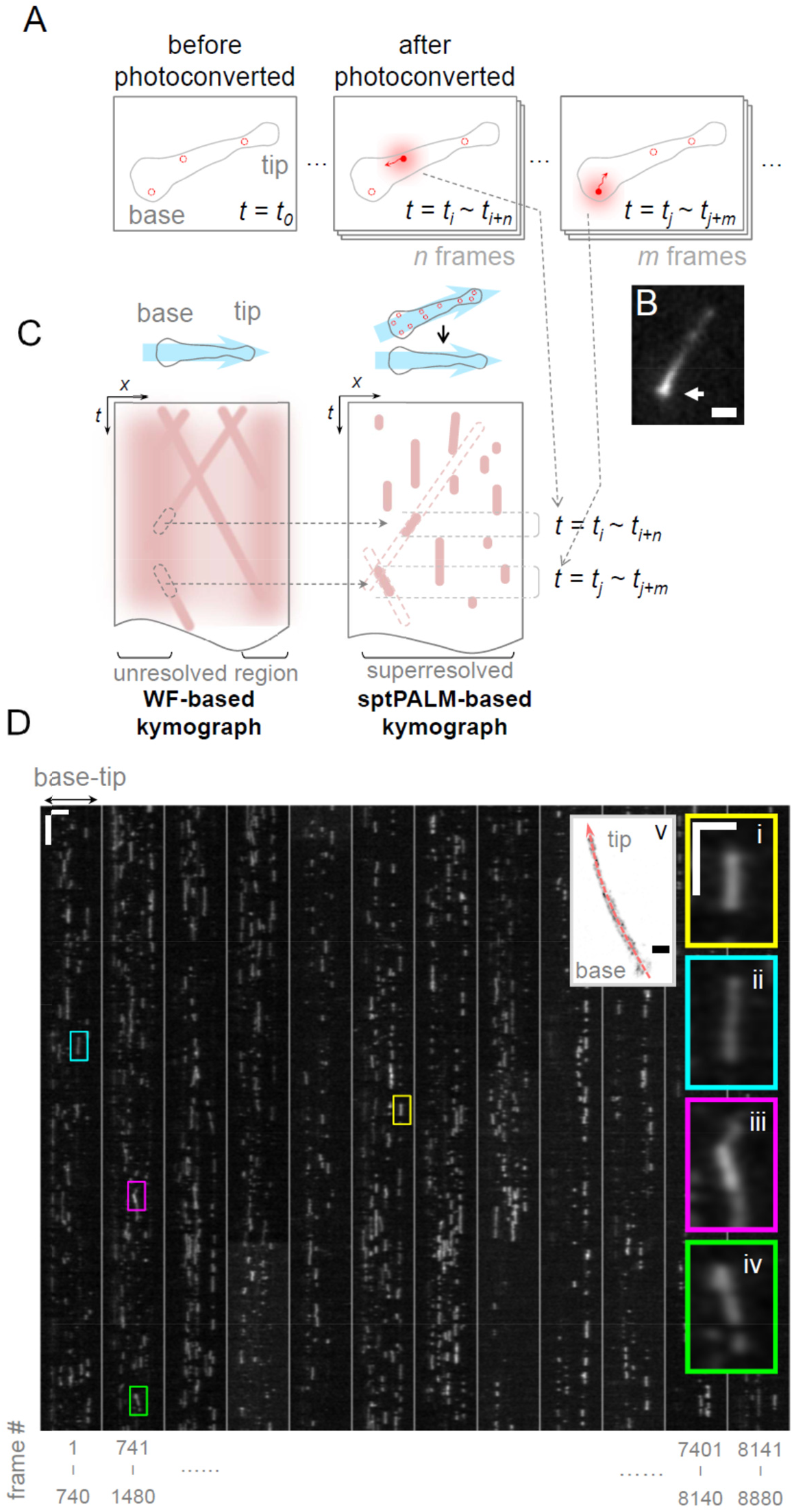
sptPALM-based imaging method for dynamic studies of IFT proteins in primary cilia. (A) Isolated single-particle tracking after photoconversion. Each probe stayed at the “on” state for several to tens of time points to create a short trajectory. (B) An epifluorescent image of IFT88-mEos4b showed a common bright punctum at the ciliary base (arrow), illustrating the high density of IFT proteins. (C) Each trajectory extracted from the defined curve along a primary cilium (blue arrow) was mapped to a trail in the kymograph (C, right); this is different from a widefield (WF)-based kymograph, which usually cannot resolve individual molecules at the ciliary base (C, left). (D) A sectioned kymograph of thousands of IFT88-mEos4b trails was reconstructed and extracted from the line shown in the live-cell PALM image (panel v). One stationary and three moving IFT proteins (observed along the axoneme) are shown in panel i and panels ii–v, respectively. Panel ii represents the retrograde motion, while the anterograde motion is shown in iii and iv. Scale bars: 1 μm [B]; vertical = 1 s and horizontal = 2 μm [D, main panel]; vertical = 500 ms and horizontal = 1 μm [D, i-iv]; 500 nm [D, v].

Thousands of trails were observed in an experimental run, where each trail represented a short trajectory of IFT88-mEos4b. Compared to a widefield-based assay where one frequently sees an ultra-bright vertical “band” of IFT proteins of a kymograph analysis present at the ciliary base (Figure S5 [7, 9, 13–15]), sptPALM enables isolation of events among the high density of IFT proteins packed in the tiny volume surrounding the distal appendages (DAPs) and transition zone (TZ) (Figure 1C). Figures 1D and S6 show sectioned experimental kymographs reconstructed from single experimental runs for a living and a fixed cell, respectively. As representative examples, one stationary molecule (or only moving laterally) and three molecules moving longitudinally are shown in panel i and panels ii—iv of Figure 1D, respectively. To analyze dynamic characteristics of IFT88 molecules from the large number of superresolved localizations, we utilized an automated, computer-based algorithm to identify each single particle position at each time point and generate trajectories from these positions (see *Methods)*. The big data from such a large number of trajectories allows us to extract statistically meaningful information from dynamic events with large variations of kinematic properties.

### The distal appendages and the transition zone accommodated not only axonemal but also transverse IFT88 movement

The recording of short superresolved trajectories for IFT88-mEos4b molecules enabled statistical analysis of their displacement directions at the ciliary base. All localizations of IFT88-mEos4b of one experimental run were overlaid as the outline of the cilium, with three trajectories illustrated showing possible paths of IFT88 at the ciliary base (Figure 2A, Movies S2-S5). Despite dynamic transport of IFT88 proteins, the image with overlaid trajectories revealed the envelope of IFT88 axonemal distribution within the ciliary compartment (CC) and the three-puncta organization at the ciliary base, agreeing with our previous finding [16, 17]. One of these example trajectories showed an IFT88 molecule moving in the transverse direction from periphery to the center of the ciliary base (Figure 2A, Movie S3), illustrating the possibility of IFT protein movement without following the DAP or axonemal structures.

**Figure 2.**
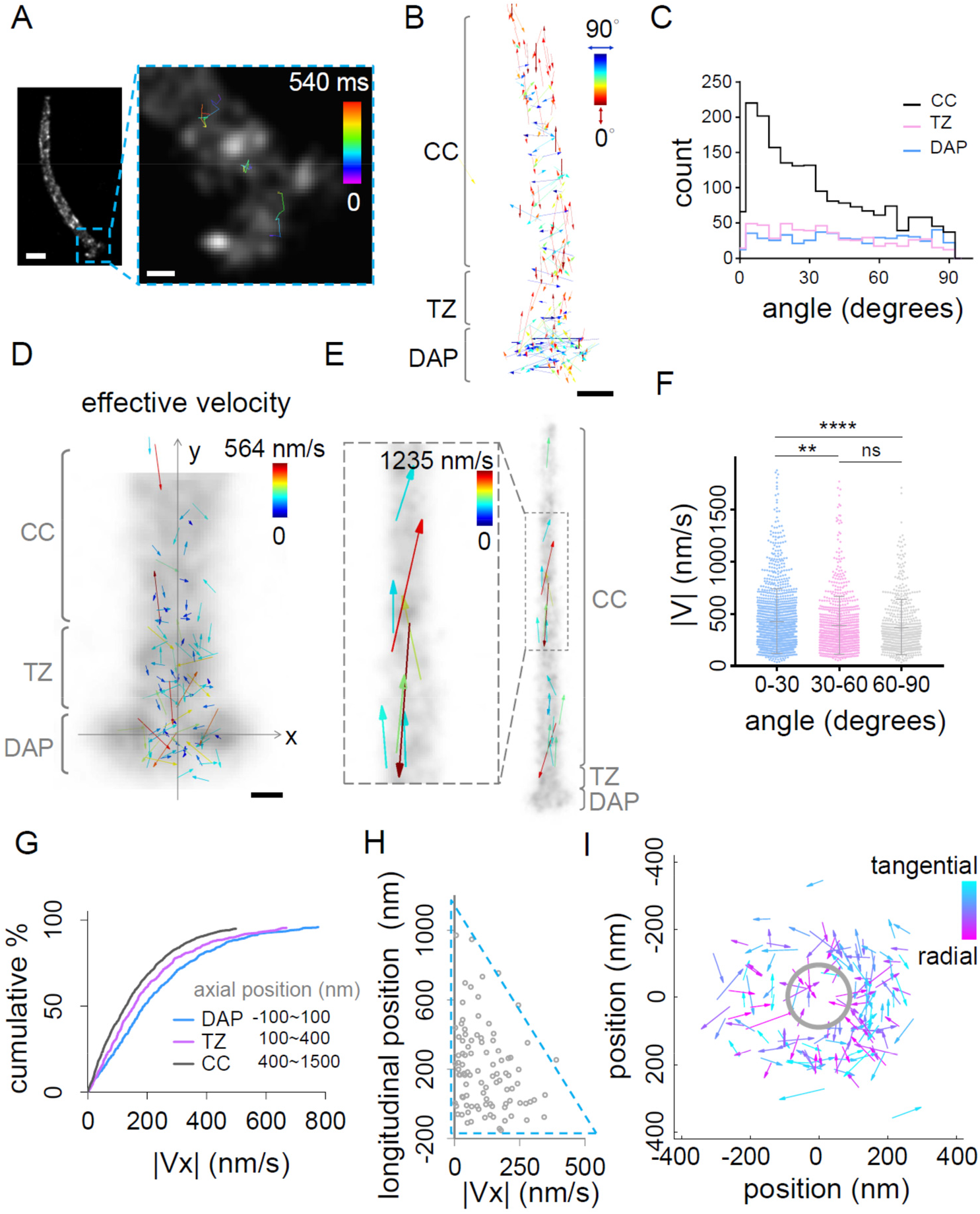
sptPALM trajectory analysis of IFT88 at the base of primary cilia showing distinct patterns of speed and direction of motion. (A) Live-cell superresolved localizations and three representative trajectories of IFT88-mEos4b. The overlaid image shows the envelope of the IFT88 axonemal distribution (left). Three trajectories of IFT88 at the ciliary base (right) show the paths of stationary and mobile IFT proteins. (B) Overlaid vectors representing effective displacements from one cilium revealed distinct angular distributions between cilium and base. The cilium (CC) had a large population of small-angle vectors, whereas the base exhibited a diverse angular distribution. (C) Detailed angular analysis from multiple primary cilia (2786 trajectories, 18 cilia) showed that the percentage of small-angle displacements (<30°) increased in the proximal–distal direction: DAPs < TZ < CC. (D) The effective velocity field calculated from the effective displacement and total travel time. (E) Representative fast-moving IFT proteins observed in the CC. (F) Statistical analysis of the orientation dependence of the effective velocity at the base were performed to compare three angular populations based on ~2800 trajectories. The result suggests that high-speed IFT movement occurred when moving approximately parallel to the axoneme. Statistics notation: ****, p < 0.0001; **, p < 0.01; ns, not significant. (G) Cumulative population plot of lateral speed (V_x_) shows that it was higher in the DAP and TZ regions than in the CC. (H) The upper bound of the V_x_ distribution decreased with longitudinal position. (I) Axial view of sptPALM-based analysis revealed that IFT88 proteins of non-ciliated cells could travel radially between the DAPs and axoneme as well as moving tangentially. Scale bars: 500 nm [A]; 100 nm [A, enlarged panels and D]; 200nm [B]. (*H*)

Other trajectories might not show clear directions of movement because of the combined effects of their short durations and stochastic motion, as well as the noise-related uncertainty in localizing single molecules (Figure 2A, Movies S4 and S5). Thus, to pinpoint the trajectories that represent directional movement, an effective displacement was defined by a simple linear regression fit of each trajectory of the mobile IFT proteins that spanned >45 nm (Figure S7). Overlaid vectors of effective displacements from multiple primary cilia (straightening and alignment procedure shown in Figures S8 and S9) showed surprising orientation difference between the cilium and the base. Most mobile IFT88 molecules within the CC moved approximately parallel to the longitudinal direction, while mobile IFT88 at the base, including the TZ and the DAPs, traveled at various angles (Figures 2B, 2C, Figure S10). Within the DAPs and TZ, we observed many translocations at near 90°, suggesting that IFT proteins might travel transversely with their nonaxonemal degrees of freedom in the space. Flux analysis of the IFT88 trajectories showed that our system approximately reached a steady state (Figure S11).

### IFT proteins had a lower transverse velocity toward the ciliary compartment

The effective velocity calculated from the effective displacement indicated that some IFT proteins moved faster than others in a location- and orientation-dependent manner (Figure 2D). Fast-traveling IFT proteins in the CC had a mean effective velocity of ~600 nm/s (Figure 2E), consistent with the velocity reported in literature [18, 19], confirming that our assay did not perturb IFT dynamics. Analyzing the orientation dependence of the effective velocity at the base, we found that high-speed IFT movement (e.g. V > 500 nm/s) occurred statistically more frequently when motion was approximately parallel to the longitudinal axis of the cilium (0/30°, Figure 2F), possibly suggesting that axoneme-associated translocation exhibited high speed but only happened occasionally. The fast-moving IFT88 at the base was slower than its fast-moving equivalents in the CC, and this may be due to the more crowded and intricate architecture of the DAP and TZ hindering movement.

To investigate the dependence of transverse velocity on longitudinal position, the lateral component of velocity (V_x_) for each moving trajectory was extracted. Consistent with our findings regarding displacement, IFT88 exhibited higher lateral speed in the DAP and TZ regions than in the CC (Figure 2G). Interestingly, IFT proteins still possessed considerable lateral speed at the TZ, suggesting that the space close to the Y-links allows lateral movement (Figure 2H). Overall, our sptPALM-based study revealed distinct patterns for IFT88 transport along primary cilia.

### Superresolved axial view revealed radial transport of IFT proteins at the DAPs

Although the lateral view showed the transverse movement of IFT proteins at the DAP region, it was unclear whether these proteins traveled along the periphery of the DAPs or radially between the DAPs and the axoneme/centriole. An axial view of IFT88 movement (here molecules were observed in non-ciliated centrioles) revealed that some IFT proteins moved tangentially to the periphery, while others moved radially. Therefore, IFT proteins could indeed travel between the DAPs and the axoneme through the space in between (Figure 2I), and the directions of travel spanned all angles, suggesting that the movement was not guided by the pinwheel-like DAP blades.

### Diffusion analysis and stepwise displacement analysis indicated that IFT protein movement was more diffusive at the proximal TZ than at the distal TZ

To further refine the spatial resolution of our dynamic characterization, we performed frame-by-frame stepwise displacement analysis for each trajectory. The map of the displacement of every two consecutive points from a trajectory showed location-dependent stepwise displacement of IFT proteins (Figure 3A). Multiple large translocation steps (e.g. those >100 nm) were detected at the proximal region of the TZ (Figure 3B), suggesting that the proximal TZ may support a fast-moving environment for IFT proteins, significantly different from the distal TZ region, where smaller translocation steps were observed (Figure 3C). The distal TZ, which is close to the location of the Y-links, might contain molecules hindering the motion of IFT proteins. Relatively small stepwise displacement was also observed at the DAP region (Figure 3C). Analyzing location-dependent consecutive displacements, we found that the regions of a higher IFT88 population density accommodated molecules that have a smaller mean consecutive displacement, and vice versa (Figure 3D), supporting the hypothesis of obstructing movement at the distal TZ and the DAP region.

**Figure 3.**
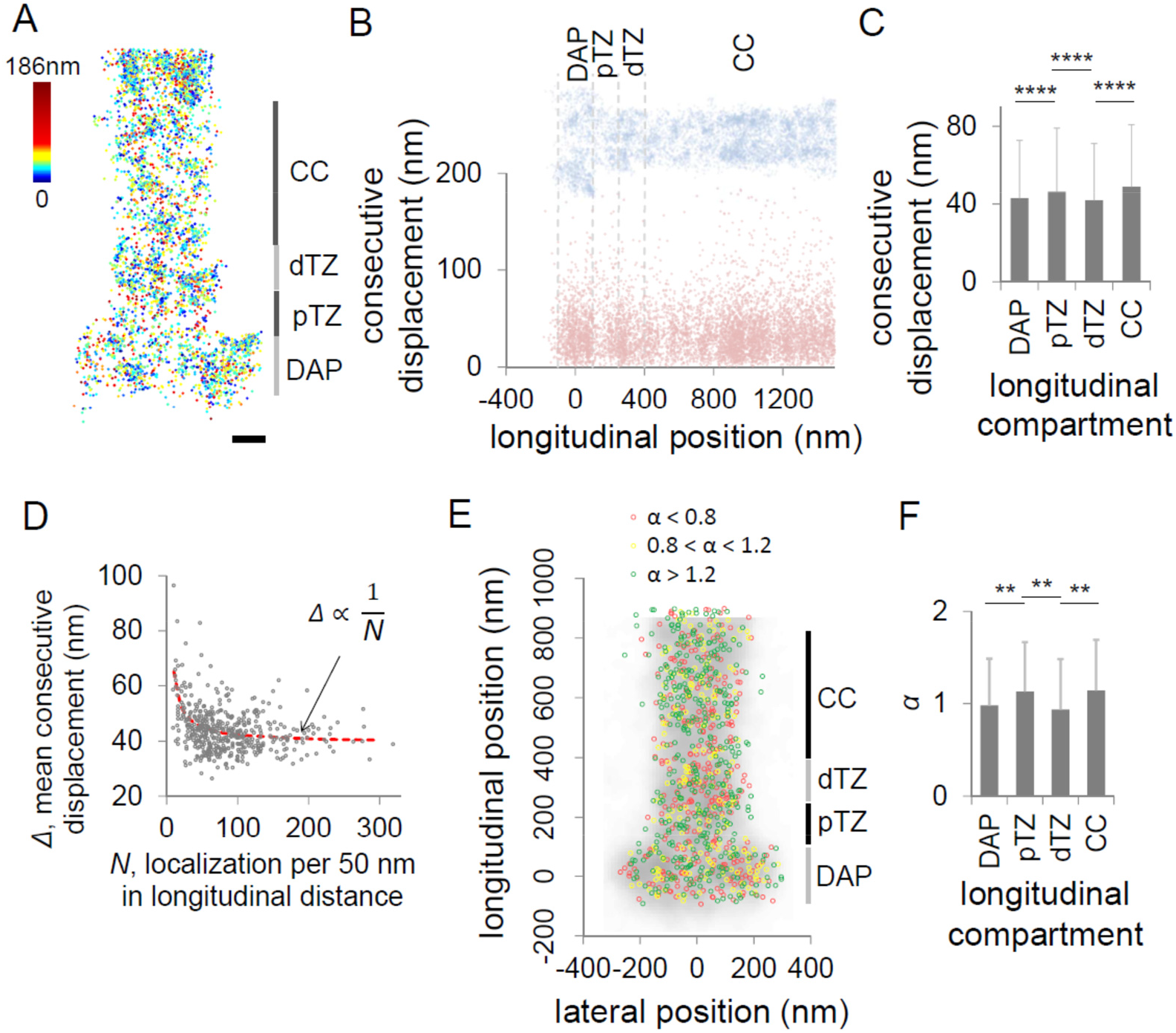
Stepwise displacement and diffusion analyses showing longitudinal dependence of IFT-protein dynamics. (A) Stepwise displacement map of IFT88 from data of three cilia displayed differential characteristics of movement between the proximal TZ (pTZ) and the distal TZ (dTZ). (B) Localization map and consecutive translocations from data in A. (C) Statistics of consecutive displacement calculated from ~18000 translocation steps of ~2800 trajectories in 18 cilia showing different translocation step sizes of the DAP, pTZ, dTZ, and CC. (D) Relation between mean local displacement and localization density illustrating that the high population regions accommodate slow moving molecules. Solid line represents the fitting result. (E) Classification of each trajectory based on diffusion characteristics *α* was mapped onto an averaged superresolved image of a primary cilium to show its location dependence. (F) Statistical analysis of longitudinal-position dependence showed that IFT protein trafficking was less diffusive at the dTZ but more diffusive at the pTZ (663 trajectories). Scale bars: 100 nm [A]. Statistics notation: ****, p < 0.0001; **, p < 0.01, Student’s *t*-test. Error bars represent standard deviation (s.d.).

Diffusion analysis further supports our observation in stepwise displacement analysis, revealing distinct diffusion mechanisms at the ciliary base (Figures 3E, 3F). We found that IFT proteins were more directionally moving in the CC, whereas in the TZ and DAPs, diffusion modes were considerably more heterogeneous. Statistical analysis of the dependence of the diffusive parameter *α* on longitudinal position showed that IFT protein movement was less diffusive at the distal TZ than in the neighboring regions (Figure 3F).

Combining the comparison of consecutive displacement and diffusion parameter, we thus concluded that IFT88 molecules change dynamic characteristics multiple times along the ciliary path, moving slowly at the DAP region, relatively fast at the proximal TZ, slowly again at the distal TZ, and then much faster in the CC.

### Two-color superresolution imaging revealed that the distal transition zone is a potential assembly site

To understand why DAPs, proximal TZ, and distal TZ had different IFT88 dynamic characteristics, we examined where IFT88 might interact with other proteins, specifically KIF3A, the kinesin responsible for anterograde movement carrying IFT proteins in the CC, and BBS2, one of the BBSome components serving as an adaptor between IFT88 and cargo proteins [20], in fixed RPE-1 cells. Two-color dSTORM superresolution imaging of BBS2 and IFT88 reveal that BBS2 molecules were populated in the CC and distal TZ, but they were considerably sparse in the proximal TZ and absent in the DAPs (Figure 4A). An image overlaying data from multiple cilia clearly illustrate differential localization of IFT88 and BBS2, with overlapping region of these two proteins primarily at the distal TZ, and some in the CC (Figure 4B). That is, the distal TZ, or possibly the CC could serve as an assembly site for IFT88 and BBS2. This result might explain why we saw the slowest motion of IFT88 at the distal TZ.

**Figure 4.**
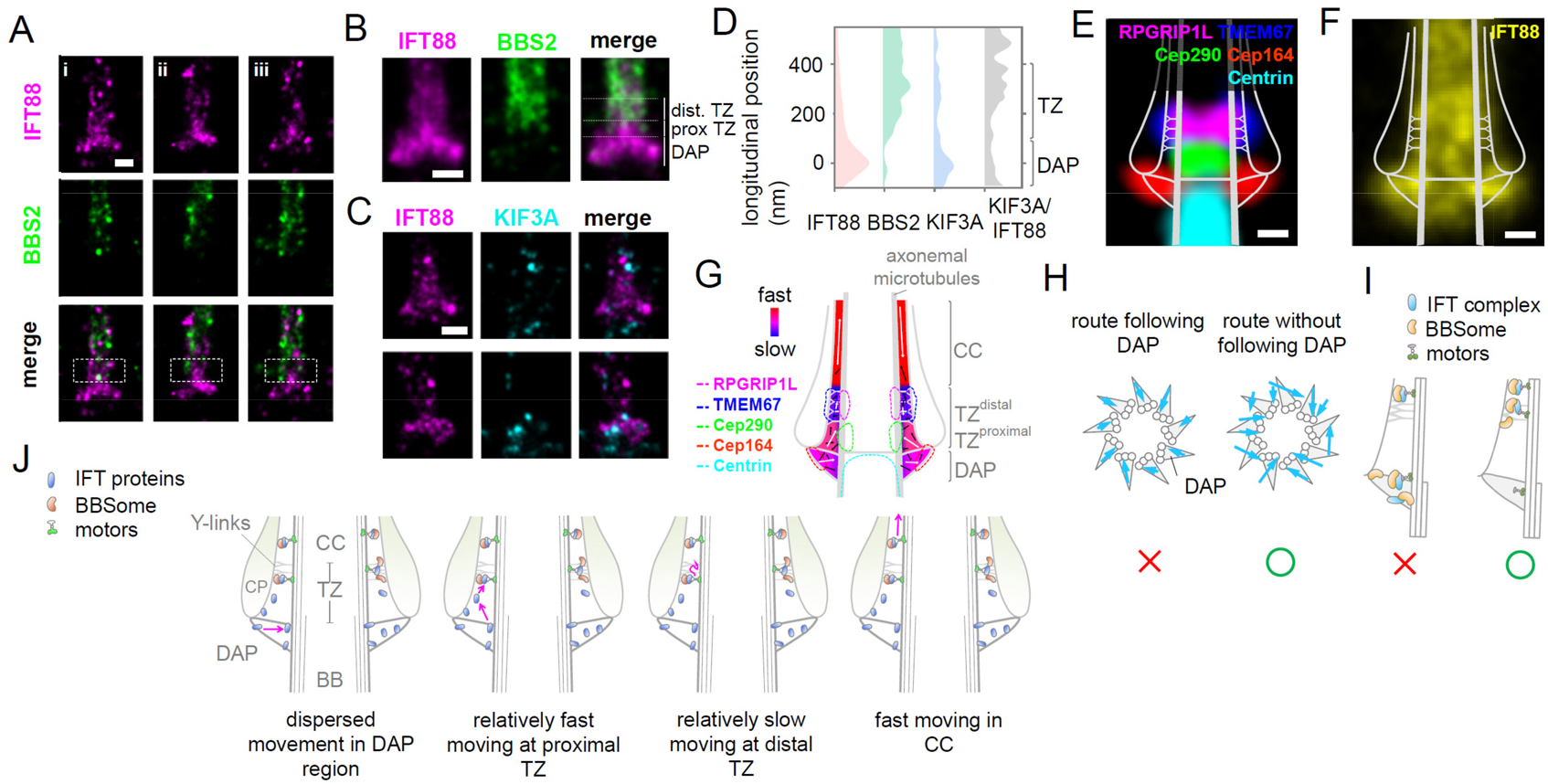
Superresolution imaging studies supporting a model of IFT protein trafficking in different regions of the ciliary base. (A) Two-color superresolution images of BBS2 and IFT88 showing that BBS2 molecules were nearly absent at the DAPs and sparsely localized at the TZ (white box), more toward the distal end. (B) Overlaid superresolution images of multiple cilia (n = 11) illustrate that BBS2 and IFT88 primarily overlapped at the distal TZ. (C) Two-color superresolution images of KIF3A and IFT88 revealing that unlike IFT88, KIF3A is less populated at the ciliary base. (D) Comparison of population distributions of IFT88, BBS2, and KIF3A quantitatively suggests their spatial coincidence for potential interactions. The ratio of KIF3A/IFT88 implies the relative probability for each IFT88 molecule to find a KIF3A molecular motor. (E) Superresolved molecular mapping of TZ/DAP proteins (RPGRIP1L, TMEM67, Cep290, Cep164, centrin) indicated TZ proteins assembled into distinct layers, for example, RPGRIP1L and TMEM67 at the distal TZ and Cep290 at the proximal TZ [17]. The distal TZ is close to the longitudinal location of the Y-links. (F) IFT88 proteins were populated at the DAPs and distal TZ. (G) A map summarizing the sptPALM results in terms of IFT-protein speed and direction. Movement of IFT proteins was slow near the distal TZ and fast in the proximal TZ. (H) Our results enable discrimination of different hypotheses. Traveling of IFT88 from the recruiting site to the axoneme does not follow the “blades” of DAPs but instead in random directions when observing in an axial view. (I) The assembly of IFT88 and BBS2 will most likely occur at the distal TZ region. This assembly is nearly impossible to happen at the DAPs because BBS2 is absent there. (J) A model illustrates the movement of an IFT protein at different regions from the DAPs to the CC: (1) dispersed and relatively slow movement in the DAP region, including inward movement from the tip of the DAPs to the axoneme; (2) relatively fast motion in the proximal TZ with a trajectory that can reach the space enveloped by the ciliary pocket; (3) slow motion at the distal TZ close to the Y-links where assembly with molecules such as BBSome proteins may occur; (4) engagement onto the microtubule track to facilitate fast movement along the axis of the CC. Scale bars: 200 nm [A-C]; 100 nm [E and F].

Two-color superresolution images of KIF3A and IFT88 showed that although we could see KIF3A molecules distributed along the cilia as expected, we found that the population of KIF3A was considerably sparse in the region where IFT88 molecules were populated (Figure 4C). That is, only a finite number of molecular motors were available to carry IFT88 molecules into the CC. Population analysis of IFT88, BBS2, and KIF3A along the longitudinal axis shows relative populations that might reflect the assembly probability in different locations (Figure 4D). As KIF3A was less populated in the ciliary base than IFT88, KIF3A should be the limiting factor for their assembly. The ratio of KIF3A/IFT88 implies that the relative probability for each IFT88 molecule to find a KIF3A molecular motor was higher toward the distal TZ.

Mapping onto the superresolved molecular architecture of the TZ and DAPs allows us to better understand location-dependent dynamic characteristics of IFT proteins on the ciliary structural framework (Figure 4E [17], F, G). Mobile IFT88 proteins travel slowly at the DAP region, possibly due to stochastic interactions with other molecules, fast at the Cep290-neighboring proximal TZ, very slow through the distal TZ close to the Y-links, potentially due to complex assembly and microtubule engagement, and very fast in the CC (Figure 4G).

### sptPALM imaging sheds lights on nonaxonemal movement of IFT88

Our studies allow us to reject some hypotheses and find support for others. One surprising finding is that traveling of IFT88 from the recruiting site to the axoneme does not follow the “blades” of DAPs (Figure 4H). Instead, IFT88 molecules move in random directions when observing in an axial view. Another finding is that IFT88 molecules can wonder in the space enveloped by the DAPs, the axoneme, the ciliary membrane, and the Y-links. Unexpectedly, they are not entirely axoneme-bound nor membrane-bound when traveling in the proximal TZ region, but stochastically moving in this volume. Finally, for complex assembly, it has been shown that various BBS proteins including BBS2 travels processively in the CC [21]. We found that the assembly of IFT88 and BBS2 most likely occur at the distal TZ region (Figure 4I). This assembly is nearly impossible to happen at the DAPs because BBS2 is absent there. Together, we propose a model summarizing different dynamic characteristics of IFT88 at the DAPs, the proximal TZ, the distal TZ, and the CC (Figure 4J).

## ACKNOWLEDGEMENTS

We thank Won-Jing Wang for sharing reagents and Chia-Lung Hsieh for comments on the manuscript. This work was supported by the Ministry of Science and Technology, Taiwan (Grant No. 103-2112-M-001-039-MY3), and the Academia Sinica Career Development Award.

